# Ordino: a visual cancer analysis tool for ranking and exploring genes, cell lines, and tissue samples

**DOI:** 10.1101/277848

**Authors:** Marc Streit, Samuel Gratzl, Holger Stitz, Andreas Wernitznig, Thomas Zichner, Christian Haslinger

**Affiliations:** Institute of Computer Graphics, Johannes Kepler University Linz, Linz, A-4040, Austria; datavisyn GmbH, Linz, A-4020, Austria; Department of Pharmacology and Translational Research, Boehringer Ingelheim RCV GmbH & Co KG, Vienna, A-1121, Austria

## Abstract

**Summary:** Ordino is a web-based analysis tool for cancer genomics that allows users to flexibly rank, filter, and explore genes, cell lines, and tissue samples based on pre-loaded data, including The Cancer Genome Atlas (TCGA), the Cancer Cell Line Encyclopedia (CCLE), and manually uploaded information. Interactive tabular data visualization that facilitates the user-driven prioritization process forms a core component of Ordino. Detail views of selected items complement the exploration. Findings can be stored, shared, and reproduced via the integrated session management.

**Availability and Implementation:** Ordino is publicly available at https://ordino.caleydoapp.org. The source code is released at https://github.com/Caleydo/ordino under the Mozilla Public License 2.0.

## 1 Introduction

A common approach in data-driven knowledge discovery is to prioritize a collection of items, such as genes, cell lines, and tissue samples, based on a rich set of experimental data and metadata. Applications include, for instance, selecting the most appropriate cell line for an experiment or identifying genes that could serve as potential drug targets or biomarkers. This can be challenging due to both heterogeneity and size of the data and the fact that multiple attributes need to be considered in combination. Advanced visual exploration tools — going beyond static spreadsheet tools, such as Microsoft Excel and Google Spreadsheets — are needed to aid this prioritization process. However, powerful general-purpose tools such as Tableau and Spotfire, on the one hand, are insufficient, as they have limitations with respect to the aggregation and visualization of genomics data. Specialized visualization tools that have proven to be valuable for exploring, ranking, and aggregating tabular genomics data (e.g., iHAT by Heinrich et al. (2012), Guided StratomeX by Streit et al. (2014)), on the other hand, have no data preloaded, lack important features due to their narrow focus, or are not (or no longer) publicly available.

To fill this gap, we present Ordino, an open-source, web-based visual analysis tool for flexible ranking, filtering, and exploring of cancer genomics data (**Fig. 1**). Furthermore, we demonstrate the use and effectiveness of Ordino in two case studies (**Supplementary Notes**, **Supplementary Figs. S2–S10**, and **Supplementary Video**).

**Fig. 1.**
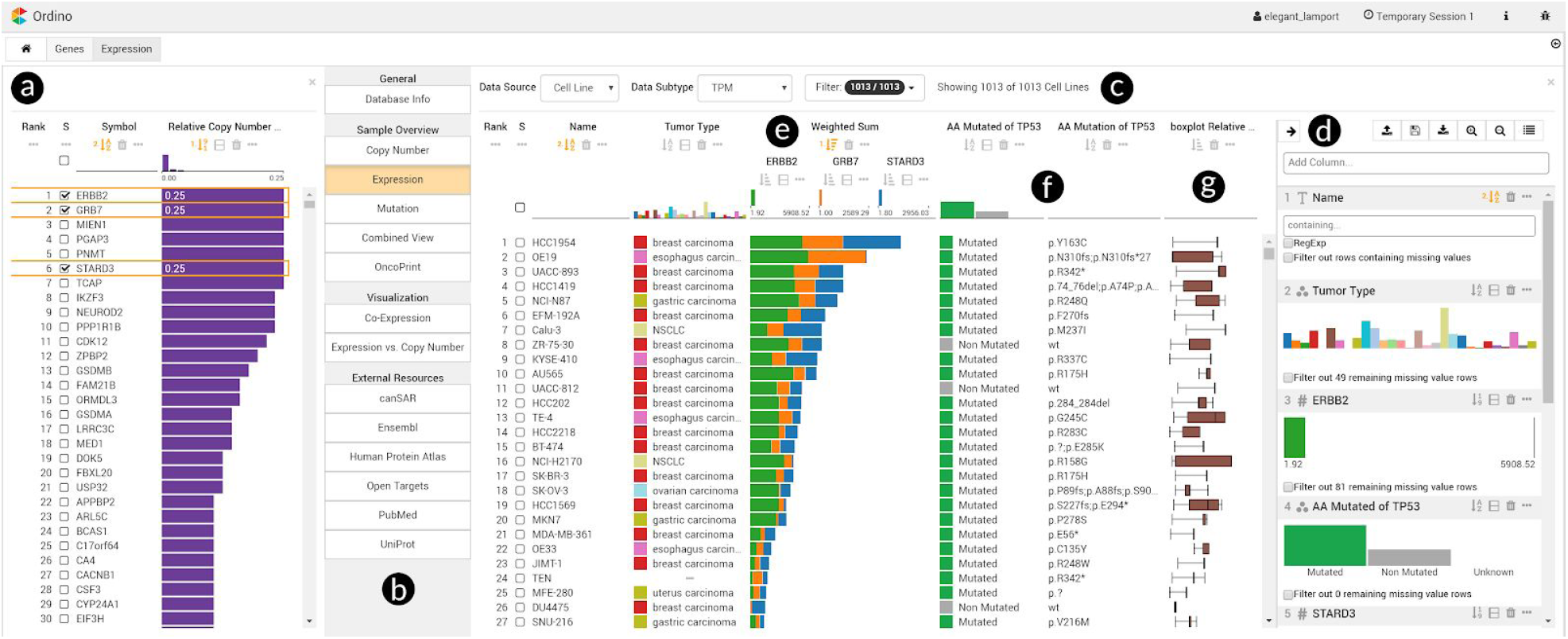
Ordino state showing genomic alteration and gene expression data of breast cancer cell lines. In the left panel (a), all human protein-coding genes are ranked by their relative amplification frequencies in a set of about 60 breast cancer cell lines. The user selects three of the most frequently amplified genes (ERBB2, GRB7, STARD3) and opens a detail view (b) on the right, displaying the expression of these genes across a set of over 1,000 cell lines (c). The side panel (d), which is shown on demand, enables the user to define a ranking hierarchy and to set filters. By combining the three gene expression columns to stacked bars (e) the user can identify cell lines in which one or more of these genes might play an important role. Next, the user adds two columns (f) that represent the mutation status and actual mutations of the cancer gene TP53. Further, a column visualizing the distribution of copy number values across 15 frequently amplified breast cancer genes is loaded (g). Based on the added information, the user gains various insights, including that the cell line with the highest expression of the three genes of interest is HCC1954, which has a p.Y163C TP53 mutation. Link to Ordino state shown in this figure: http://vistories.org/ordino-teaser-figure

## 2 Software Description

Ordino enables users to flexibly prioritize and explore items based on a rich set of experimental data and metadata. The basic workflow comprises three steps:

1. Select or define a list of items consisting of either genes, cell lines, or tissue samples, which determines the rows in a tabular visualization. For this list of items the user can add or upload multi-attribute data that is shown as columns in the table.
2. Prioritize, filter, and explore the items using an extended version of our interactive ranking technique LineUp (http://lineup.caleydo.org; Gratzl et al., 2013) (**Fig. 1a** and **1c**, **Fig. S3**).
3. Obtain detailed information about one or multiple items of interest by selecting them in the table and opening various detail views.

### 2.1 Item List Definition and Adding Data

In Ordino the user starts the prioritization by defining a set of items. The item set can be determined by manually entering a list of identifiers (e.g., a list of gene symbols or cell line names), by selecting a previously saved or predefined list of items, or by uploading a comma-separated file (**Fig. S2**).

Subsequently, the user can interactively add (i) raw experimental data or metadata stored in the Ordino database, such as the expression data for a single cell line or the biotype of all listed genes, (ii) dynamically computed scores, such as the average gene expression of tissue samples from a specific tumor type, and (iii) uploaded custom data attributes, which allows the user to fuse external data to the currently shown table.

We preloaded the Ordino database with mRNA expression, DNA copy number, and mutation data from The Cancer Genome Atlas (TCGA; https://cancergenome.nih.gov) and the Cancer Cell Line Encyclopedia (CCLE; Barretina et al., 2012), as well as two depletion screen data sets from McDonald et al. (2017) and Meyers et al. (2017) (**Supplementary Table S1**). A description of the data pre-processing can be found in the supplementary material (**Supplementary Notes**).

### 2.2 Interactive Visualization of Rankings

The tabular data is visualized using an extended version of our interactive ranking technique LineUp. Users can change the visual representation of columns on demand. Single numerical attributes can be visualized, for instance, by using bars, by varying the brightness, or by means of circles whose sizes are proportional to the data values. Columns containing dynamically computed scores that aggregate data from multiple entities can be shown as row-wise box plots or heat maps visualizing the raw data values that constitute the aggregated score. Users can change the visual representation of columns on demand. The exploration is supplemented with filtering features such as setting cut-off values for numerical columns and specifying one or multiple categories in categorical columns.

Furthermore, users can rank the table by a single column (e.g., the value of a numerical column or the median of a box-plot column) or by interactively created weighted combinations of two or more columns. The combined column is then shown as a stacked bar highlighting the contribution of individual attributes to the total score. More advanced combinations can be defined interactively or via a scripting interface. Examples of such advanced combinations are (i) min/mean/max combination columns, which show only the minimum, mean, or maximum of all combined columns, (ii) scripted columns, for which users can define how individual columns are to be combined using JavaScript, (iii) nested columns for semantically grouping multiple columns, and (iv) imposition columns, which color numerical columns by a categorical attribute.

### 2.3 Detail Views

Users can select one or multiple items in the table to explore them using a collection of detail views (**Fig. 1b**, **Supplementary Notes**). Detail views can be (i) specialized visualizations (e.g., a co-expression plot for comparing multiple genes, an expression vs. copy number plot, or an OncoPrint), (ii) another ranked table (e.g., a list of all tissue samples plus their expression, copy number, and mutation data for the selected genes), or (iii) embedded external resources (Ensembl, Open Targets, etc.). Newly opened detail views appear on the right side of the interface, causing the previously active view to be shown in a more compact format on the left.

### 2.4 Reproducibility and Sharing

A core feature of Ordino is its ability to let users store, revisit, and share findings at any time during analysis sessions. To achieve this, Ordino requires users to log in before using the system. To avoid a tedious registration process for creating the login credentials, the system auto-generates temporary accounts. A session keeps track of all interactions and data uploaded by the user throughout the analysis. By default, analysis sessions are temporary, which means that they are stored only in the local cache of the browser. The user can choose to make a session persistent, which moves it from the local browser storage to the database on the Ordino server. Persistent sessions can be shared by simply copying the URL shown in the browser. When a link to a persistent session is opened, the system restores the exact state of the analysis, including the history of all previous steps (Gratzl et al., 2016). Note that the states shown in **Fig. 1** and **Supplementary Figs. S2-S10** can be reproduced by following the links provided below the figures.

## 3 Implementation

The Ordino software is based on the extensible Phovea platform (http://phovea.caleydo.org). The web client is implemented in TypeScript, and the server in Python. The public version of the system is deployed on Amazon Web Services (AWS) infrastructure using Docker images (https://www.docker.com). The source code together with more information on how to install Ordino locally and how to add custom data connectors that fetch data from public REST APIs or from relational databases can be found on Github (https://github.com/Caleydo/ordino).

## 4 Conclusion

We believe that Ordino is a powerful cancer genomics tool for flexibly prioritizing and exploring items based on a rich set of experimental data and metadata. It can be used for numerous purposes, for instance, to identify suitable cell lines for an experiment, investigate gene signatures, and identify and prioritize potential biomarkers. Since custom data can be loaded into Ordino, the tool can in principle also be applied in fields beyond cancer genomics. Thus, Ordino supports a wide range of users, including bioinformaticians, biologists, and researchers from other domains.

## Acknowledgements

We thank Christian Lehner for contributions to the implementation of the tool as well as Daniel Gerlach, Markus Bauer, and Anita Steiner for their contributions to data preparation and data handling.

