## Supplementary Notes for "Ordino: a visual cancer analysis tool for ranking and exploring genes, cell lines, and tissue samples"

---

---

#### Table of Contents

Supplementary Figures 2
Supplementary Tables 12
Supplementary Notes 13
Supplementary References 25

#### Supplementary Figures

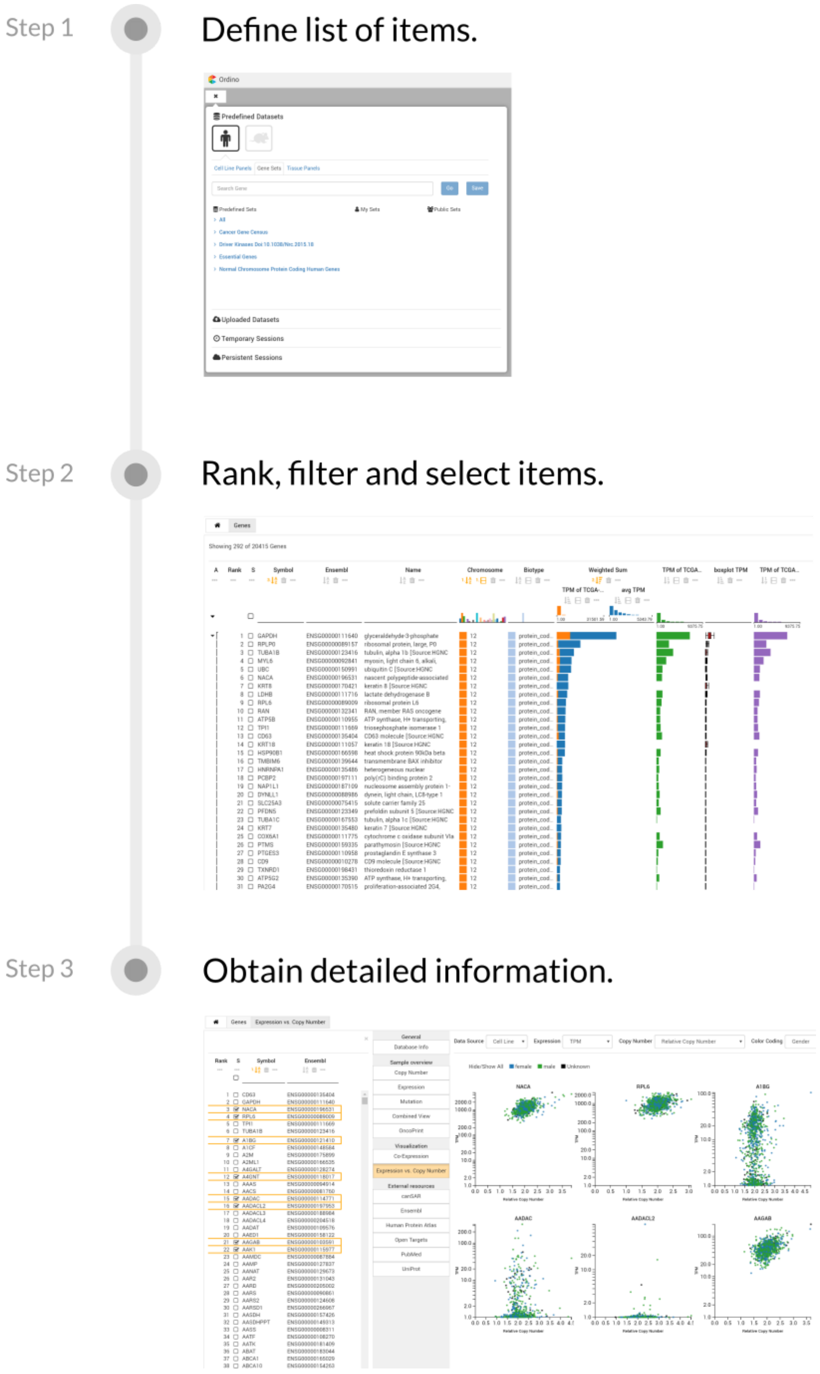


**Supplementary Figure S1. Ordino workflow.** The workflow comprises three steps. (1) Select or define a list of items consisting of genes, cell lines, or tissue samples. (2) Rank, filter, and select items. (3) Obtain detailed information for selected items.

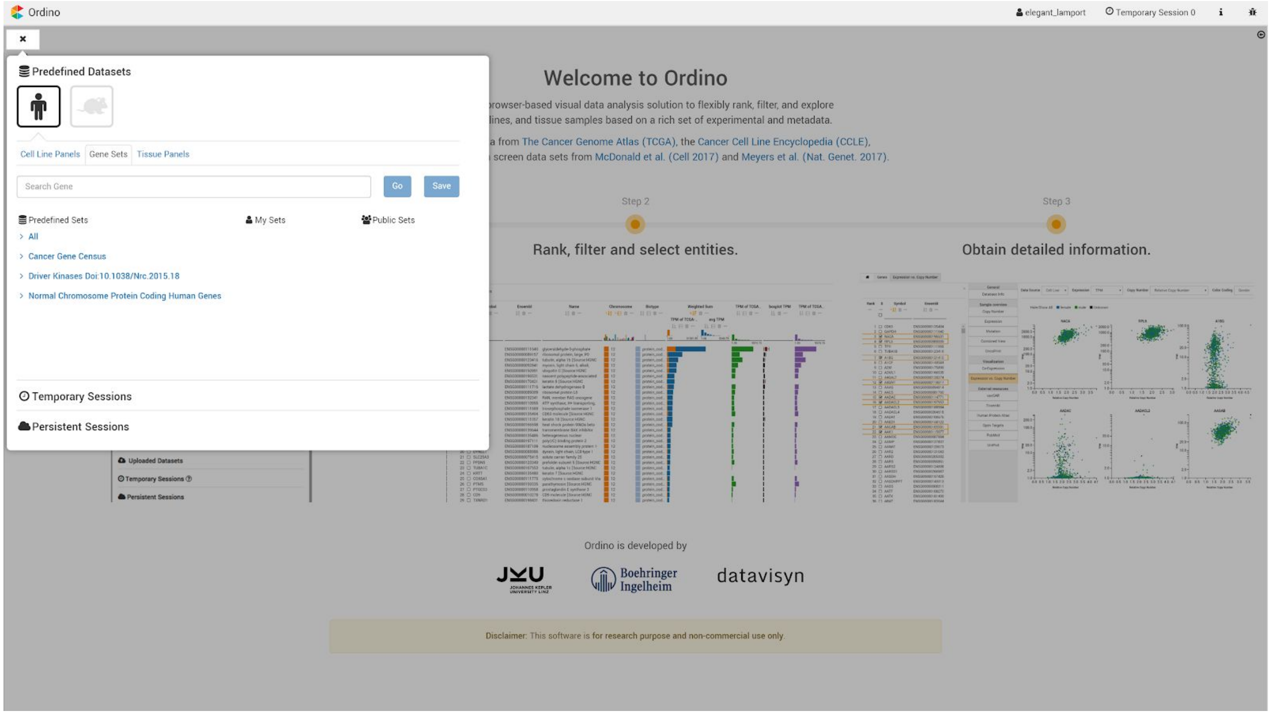


**Supplementary Figure S2. Case study 1: Definition of gene list.** In the start menu the user can choose predefined sets or previously saved public and private sets as a starting point for the analysis. Additionally, one can upload a custom dataset or continue a previous analysis session.

Link to Ordino state shown in this figure: http://vistories.org/ordino-supplementary-figure-2

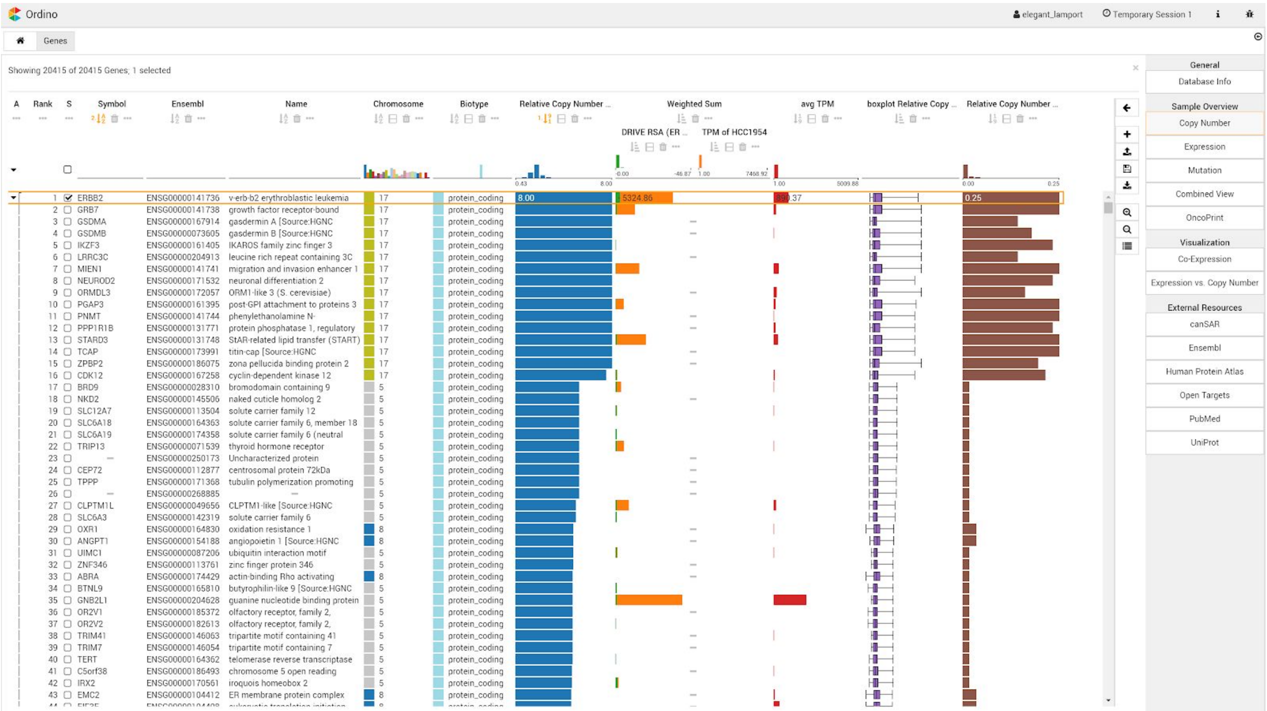


**Supplementary Figure S3. Case study 1: Ranking of genes.** Gene list with additional score columns, which are calculated on the fly. The ranking shows the highest amplification in the cell line HCC1954, located on chromosome 17, affecting about 15 genes, with *ERBB2* (*HER2*) having the highest expression level (orange column) and the lowest sensitivity score (green column). Therefore, it is probably the most relevant gene of this amplicon. The two aggregated score columns (in red and brown) show that *ERBB2* is amplified in almost 25% of all assessed breast cancer cell lines. Further, it is highly expressed.

Link to Ordino state shown in this figure: http://vistories.org/ordino-supplementary-figure-3

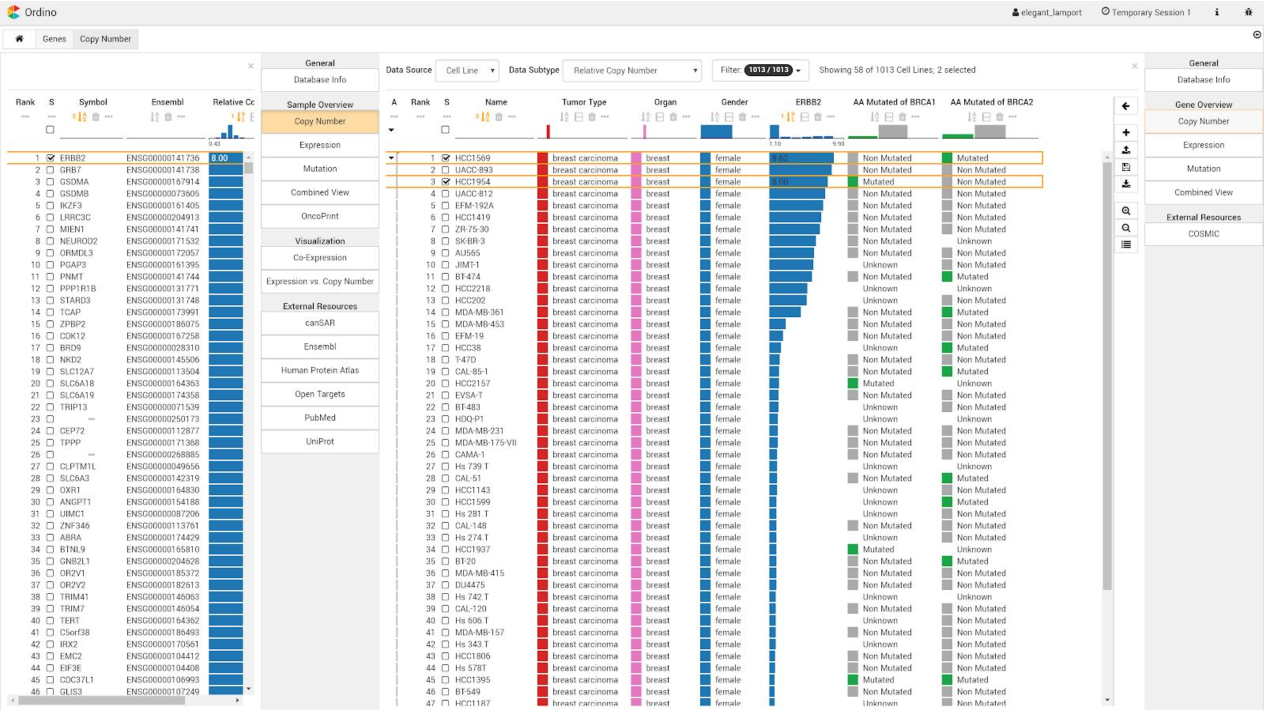


**Supplementary Figure S4. Case study 1: Cell lines ranked by copy number data of *ERBB2*.** The user selects the gene *ERBB2* and opens the Copy Number detail view as new focus view. The gene list remains open on the left as context view. The Copy Number view shows that HCC1954 has the highest *ERBB2* amplification among *BRCA1* mutated cell lines and that HCC1569 has the highest *ERBB2* amplification among *BRCA2* mutated cell lines.

Link to Ordino state shown in this figure: http://vistories.org/ordino-supplementary-figure-4

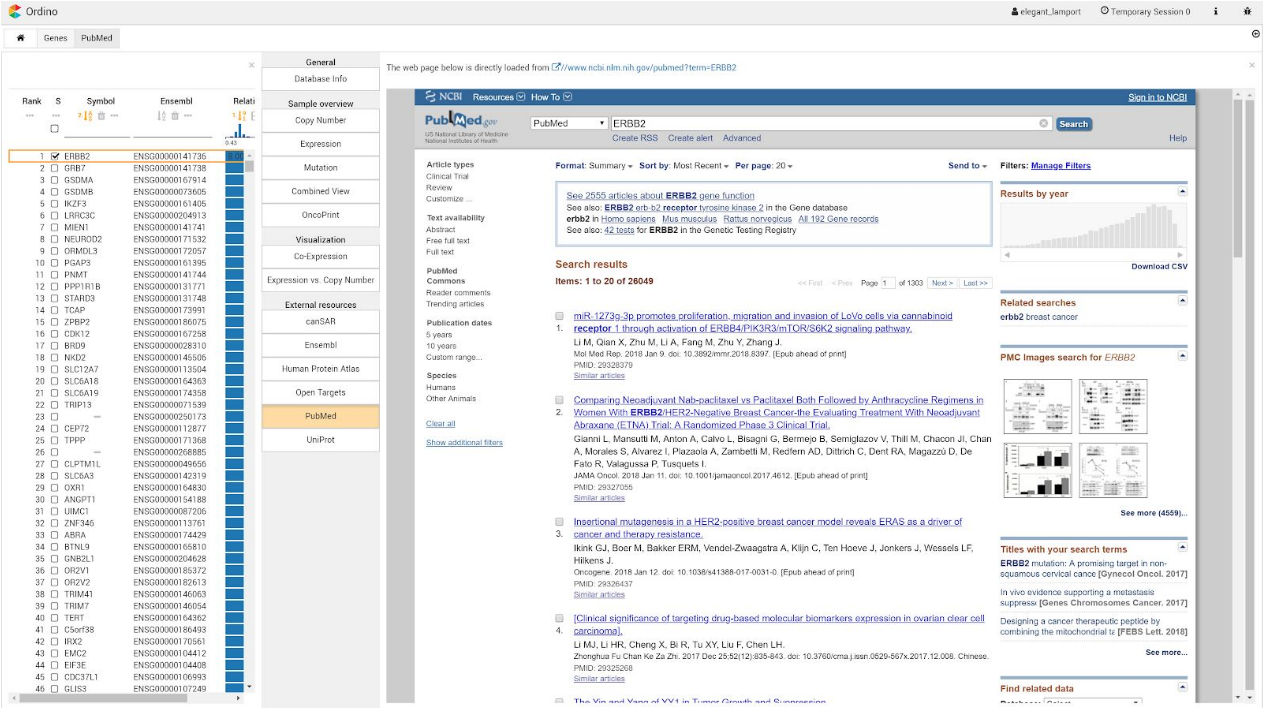

**Supplementary Figure S5. Case study 1: PubMed information for *ERBB2*.** Users can obtain further information for selected items in detail views that load the content from external websites, such as PubMed and Open Targets.

Link to Ordino state shown in this figure: http://vistories.org/ordino-supplementary-figure-5

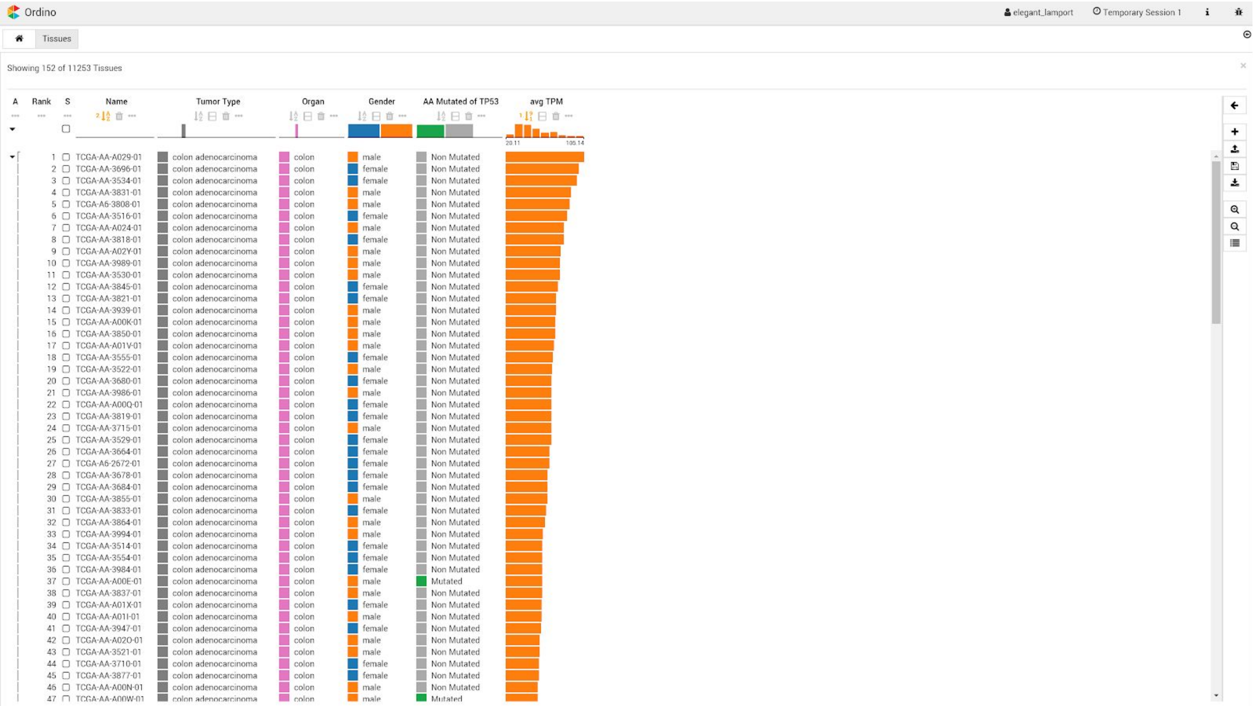


**Supplementary Figure S6. Case study 2: TCGA tumor samples ranked by average expression.** The list of TCGA tumor samples shows a clear correlation between the gene expression signature (orange bars) and the mutation status of *TP53*: Of the 50 samples with the highest expression only 3 are *TP53* mutated, whereas of the 50 samples with the lowest expression 35 are TP53 mutated (not visible in this figure).

Link to Ordino state shown in this figure: http://vistories.org/ordino-supplementary-figure-6

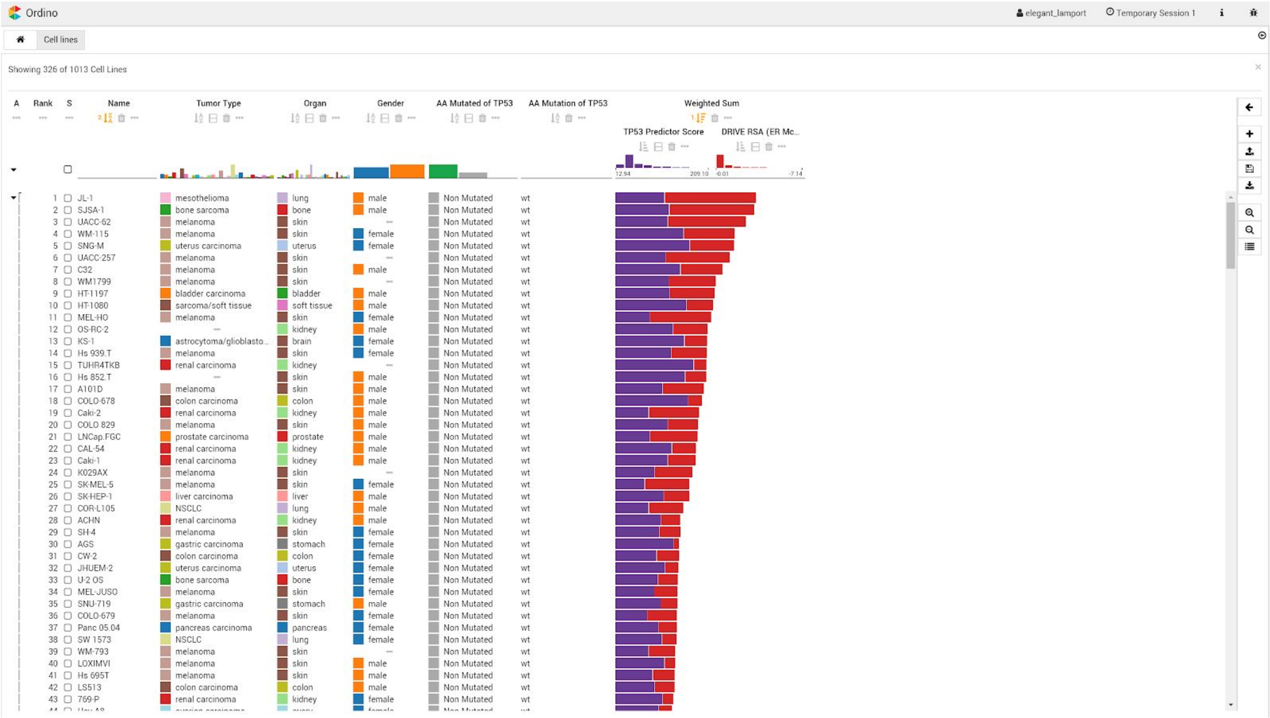


**Supplementary Figure S7. Case study 2: Cell lines ranked by weighted sum of two scores.** The analyst has added mutation and score columns to the list of cell lines. The combined score of DRIVE RSA values for the gene *MDM2* (red bars) and the gene expression signature (*TP53* Predictor Score; purple bars) correlates with the *TP53* mutation status (column AA Mutated).

Link to Ordino state shown in this figure: [http://vistories.org/ordino-supplementary-figure-7](http://vistories.org/ordino-supplementary-figure-7)

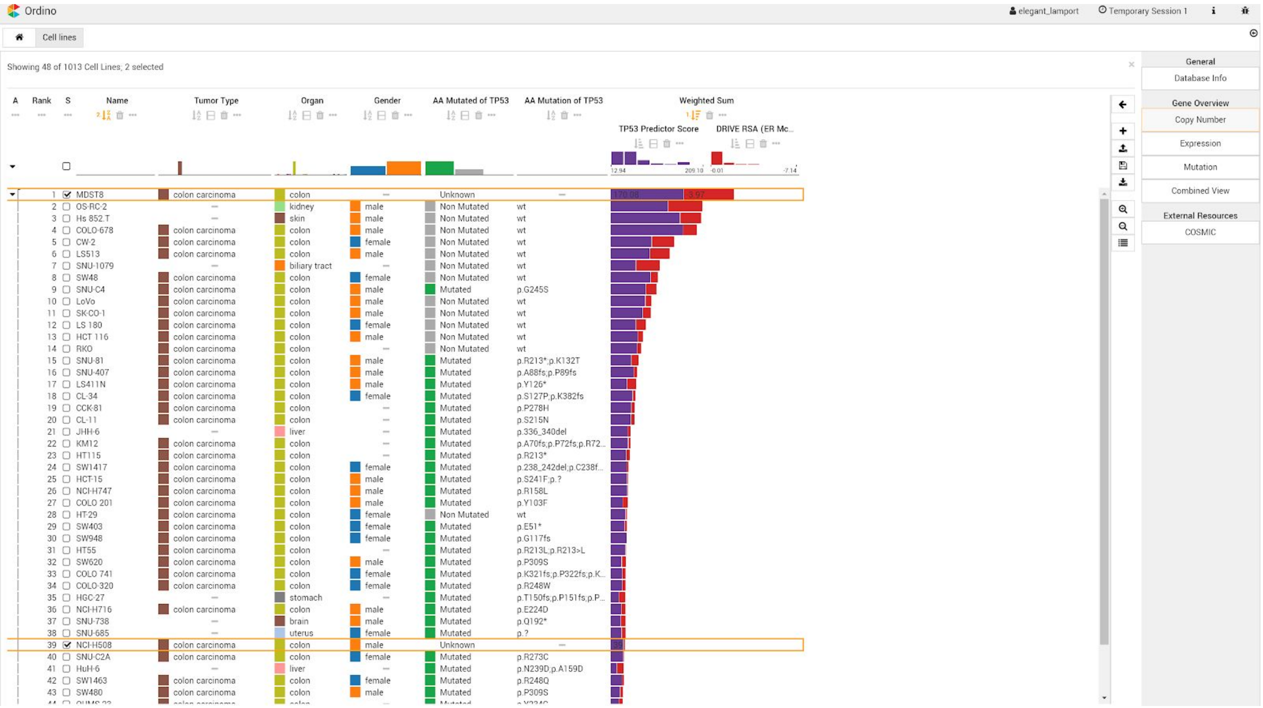


**Supplementary Figure S8. Case study 2: Filtering list of cell lines ranked by weighted sum of two scores.** Filtering the list of cell lines for tumor type *colon carcinoma* reveals that the cell lines MDST8 and NCI-H508 lack *TP53* mutation status, but gene expression and DRIVE data are available. MDST8 has a very high combined score and is therefore unlikely to be *TP53* mutated. NCI-H508, however, has a very low combined score and is therefore probably *TP53* mutated.

Link to Ordino state shown in this figure: http://vistories.org/ordino-supplementary-figure-8

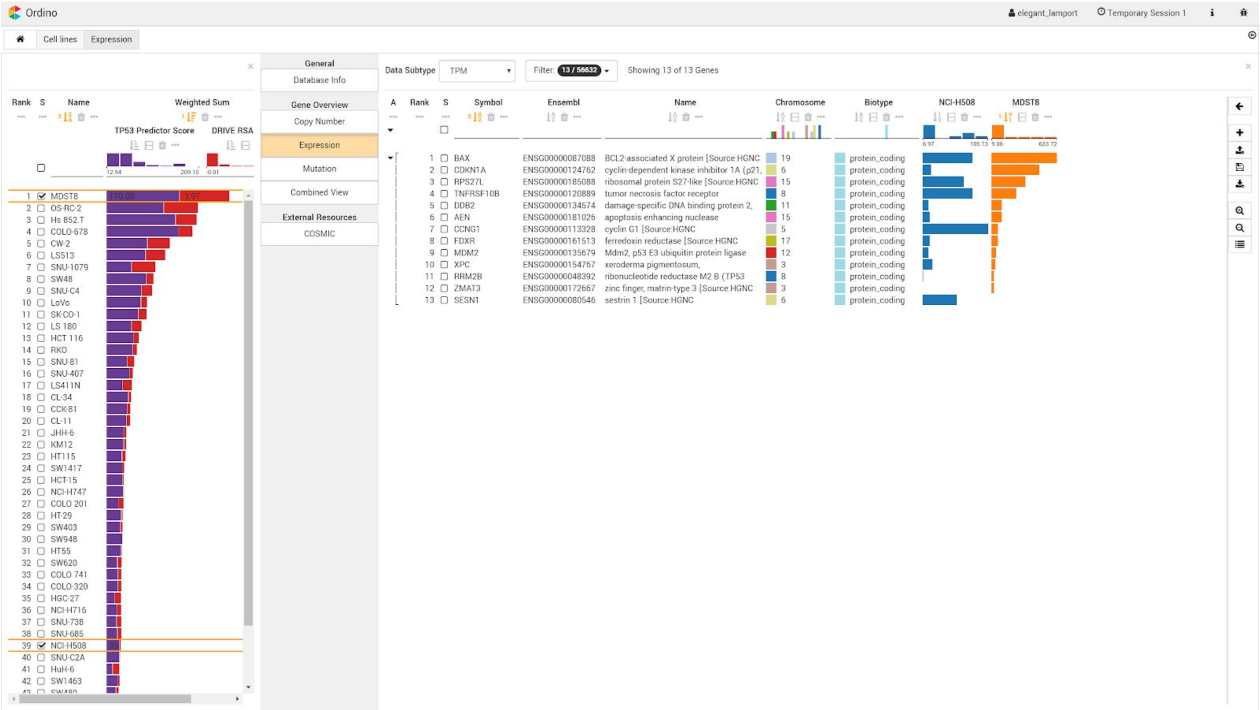

**Supplementary Figure S9. Case study 2: Gene expression detail view showing expression of two genes.** Opening the Gene Expression detail view shows that the high gene signature expression in the cell line MDST8 is caused mainly by expression of the genes *BAX*, *CDKN1A*, and *RPS27L*.

Link to Ordino state shown in this figure: [http://vistories.org/ordino-supplementary-figure-9](http://vistories.org/ordino-supplementary-figure-9)

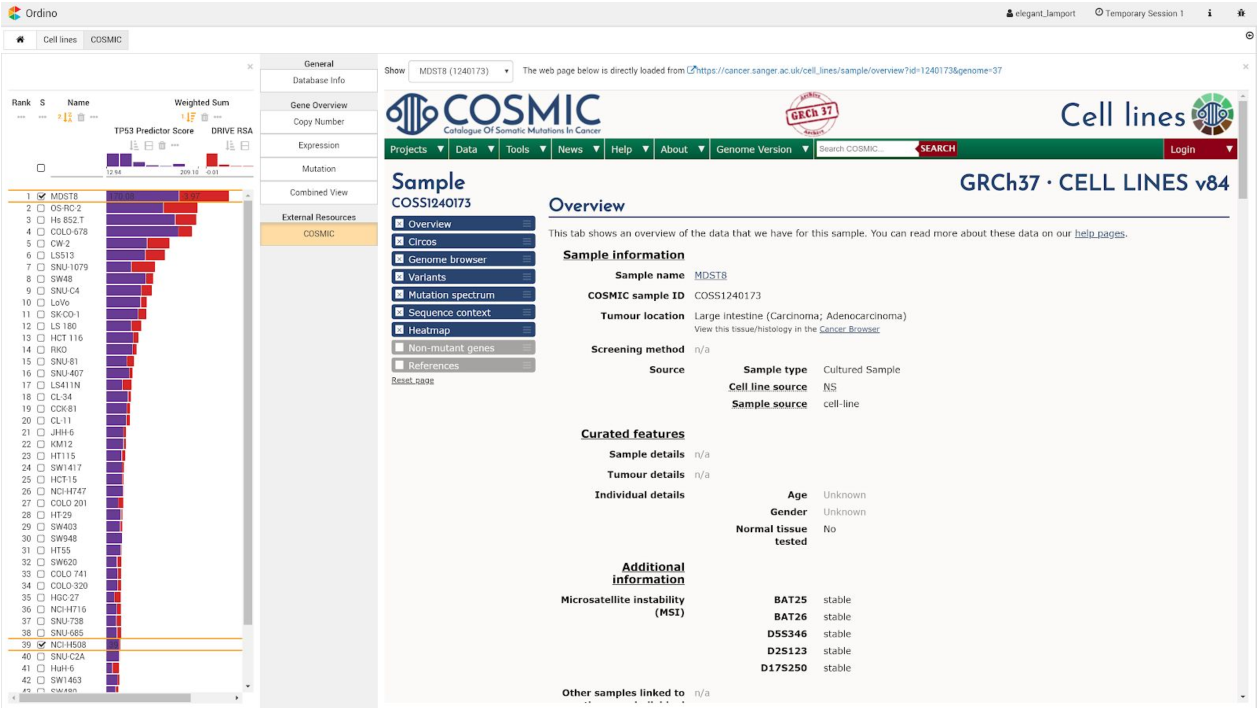

**Supplementary Figure S10. Case study 2: COSMIC detail view.** The analyst can browse the information available for selected cell lines in a detail view, in this case the corresponding COSMIC page for MDST8.

Link to Ordino state shown in this figure: http://vistories.org/ordino-supplementary-figure-10

### Supplementary Tables

**Supplementary Table S1.** Data available for cell lines and tissue samples, with mRNA expression, copy number, and mutation status data as well as number of available genes on canonical chromosomes. The gene annotation is based on Ensembl Version 70.

| *Dataset* | *Assay* | *Samples* | *Genes* | *Type* |
|---|---|---|---|---|
| tissues | | 12,334 | | |
| tissues | mRNA-seq expression | 10,623 | 19,959 | continuous |
| tissues | SNP6-array based copy number data | 12,035 | 56,595 | categorical and continuous |
| tissues | DNA sequencing based mutation data | 6,795 | 20,136 | categorical |
| cell lines | | 1,013 | | |
| cell lines | mRNA-seq expression | 932 | 56,495 | continuous |
| cell lines | SNP6-array based copy number data | 974 | 56,595 | categorical and continuous |
| cell lines | DNA sequencing based mutation data | 810 | up to 20,197 | categorical and continuous |
| cell lines | RNAi depletion screen data (Project DRIVE [4]) | 385 | 7,267 | continuous |
| cell lines | CRISPR-Cas9 depletion screen data (Avana CERES [5]) | 330 | 17,586 | continuous |
| genes | | | 56,632 | |

### Supplementary Notes

#### Background

To address the task of prioritizing a collection of items based on a rich set of experimental data and metadata, domain experts use either programming languages, such as R and Python, or tools, such as Microsoft Excel, Spotfire, and Tableau.

Programming languages provide the full flexibility needed for analysis. However, they require substantial programming knowledge, and data integration and preparation are time-consuming; even minor changes in the analysis can lead to major changes in the code. Furthermore, the output of scripts based on languages such as R and Python is usually static, which severely limits the exploratory nature of the analysis.

Powerful general-purpose tools such as Tableau and Spotfire can be too difficult to use for a non-expert and have limitations with respect to the aggregation and visualization of genomics data.

Spreadsheet tools, such as Microsoft Excel and Google Spreadsheets, are not designed for handling larger genomics datasets. Further, neither of these allows detailed information about individual cell lines or genes to be retrieved in an interactive manner.

#### Visual Analysis Workflow

The Ordino analysis workflow consists of three steps, as outlined in **Supplementary Fig. S1**.

##### *Step 1: Define List of Items*

The user starts the analysis by defining a set of items. The item set can be determined by manually entering a list of identifiers (e.g., a list of gene symbols), by selecting a previously saved or predefined list of items, or by uploading a comma-separated file (**Supplementary Fig. S2**).

##### *Step 2: Rank, Filter, and Select Items*

A core component of the Ordino system is the interactive visualization technique LineUp (http://lineup.caleydo.org) [1], which allows users to flexibly create and explore rankings of items based on a set of heterogeneous attributes. The exploration is supplemented with filtering features, such as setting cutoff values for numerical attributes, specifying a string or regular expression for textual columns, and specifying one or more categories in categorical attributes. In addition, users can change the visual representation of columns on demand. Numerical attributes, for instance, can be visualized using bars, varying brightness, or as circles whose sizes are proportional to the data values.

As a starting point, Ordino presents the list of items selected in Step 1 as a table containing metadata attributes specific to the item type. For genes, the default columns are gene symbol, Ensembl ID, chromosome, and biotype (**Supplementary Figure S3**). For cell lines and tissue samples, the default columns are name, tumor type, organ, and gender (**Supplementary Figure S6**). Initially, gene lists are sorted alphabetically by gene symbol, and lists of cell lines and tissue samples by their name. Further columns can be added by clicking on the plus icon shown on the right-hand side of the interface (**Supplementary Fig. S3**).

Ordino supports the following column types:

- **Database columns** contain metadata about genes (such as biotype, chromosome, Ensembl ID, name, sequence region start & end, strand, and gene symbol), cell lines (age at surgery, gender, growth type, histology type, metastatic site, morphology, name, organ, and tumor type), and tissue samples (age, body mass index (BMI), days to death, days to last follow up, ethnicity, gender, height, name, organ, race, tumor type, tumor type adjacent, vendor name, vital status, and weight).
- **Parameterized Score columns.** Depending on the item type of the main table, users can add single score columns by specifying a single item (gene, cell line, or tissue sample) together with the data attribute of interest (e.g., expression, copy number, or mutation). The following single scores are available:
  - For **genes**: single tissue-sample score, single cell-line score, and single depletion-screen score.
  - For **cell lines**: single gene score and single depletion-screen score.
  - For **tissue samples**: single gene score.

  In addition to single score columns, the values of which are loaded directly from the the Ordino database, users can define aggregations of multiple items that are calculated on the fly by the Ordino server. To define an aggregation, users must specify (1) the set of items on whose basis the aggregation will be calculated (either by selecting previously stored named sets, by entering lists of items, or by selecting categorical attributes, such as the tumor type of cell lines), (2) the data type (expression, copy number, mutation, and depletion screen), and (3) the aggregation function (average, median, min, max, box plot, frequency, and count). The following aggregated scores are available:
  - For **genes**: aggregated tissue-sample score, aggregated cell-line score, and aggregated depletion-screen score.
  - For **cell lines**: aggregated gene score and aggregated depletion-screen score.
  - For **tissues samples**: aggregated gene score.
- **Combining columns** allow users to combine the content of multiple columns by dragging the header of single columns onto the combined column header. Users can create **weighted sum columns**, which are visualized as stacked bars highlighting the contribution of individual attributes to the total score, **min/mean/max combination columns**, which show only the minimum, mean or maximum of all combined columns, **scripted columns**, for which users can define how individual columns are to be combined using JavaScript, **nested columns** for semantically grouping multiple columns, and **imposition columns**, which color numerical columns by a categorical attribute.
- **Uploaded columns** allow users to fuse external data to the currently shown table. The data can be loaded from a comma-separated file in which the first column contains the unique identifier of the primary identifier in the table, followed by one or multiple columns holding the data to be integrated. The system automatically detects common annotations, such as gene symbols and Ensembl IDs.

##### *Step 3: Obtain Detailed Information*

Users can select one or more items in a ranking table for exploration using a collection of detail views. The detail views offered to the user depend on the type of items selected. The publicly deployed version of Ordino includes the following detail views:

- **Database Info view** for showing metadata stored in the database for all selected items. The information is represented as a table containing a row for each database attribute and a column for each selected item.
- **Expression view, Copy Number view, and Mutation view** visualizing experimental data for the currently selected items with the ranking visualization technique described in Step 2 of the analysis workflow.
- **Combined view** is a specialized ranking view that is able to show copy number, expression, and mutation data in combination.
- **Expression vs. Copy Number view** showing a scatterplot for each selected gene with copy number mapped to the x-axis and expression to the y-axis (cf. Step 3 in **Supplementary Fig. S1**). The analyst can determine via a drop-down list whether the scatterplot shows cell lines or tissue samples. Dots in the plots can be colored by preloaded categorical attributes, such as tumor type, gender and organ.
- **Co-Expression view** for comparing the expression of multiple selected genes. If multiple genes are selected, one plot is shown for each combination. Analogously to the Expression vs. Copy Number view, the dots represent either cell lines or tissue samples and can be colored by categorical attributes.
- **OncoPrint view** showing a horizontal series of colored blocks (glyphs) for each gene selected. Depending on the chosen data subset, each block represents a cell line or tissue sample. The background color of the blocks indicates the copy number status (pink=amplification, blue=deep deletion, gray=normal, white=unknown), while the small block contained visualizes the mutation status (green=mutated, gray=non mutated, white=unknown) of a cell line or tissue sample.
- **External resource views** loading the content of external websites. For selected genes, the user can look at the information available on canSAR, Ensembl, Human Protein Atlas, Open Targets, PubMed, and UniProt. For cell lines, the analyst can load the information available on COSMIC (Catalogue Of Somatic Mutations In Cancer).

#### Implementation and Availability

Ordino is publicly available at https://ordino.caleydoapp.org and best viewed in Google Chrome browsers. The source code is freely available at https://github.com/Caleydo/ordino.

Ordino is based on the extensible Phovea platform (http://phovea.caleydo.org). The web client is implemented in TypeScript and the server in Python. The source code is open source under the Mozilla Public License (MPL) and hosted on Github. The public version of the system is deployed on AWS (Amazon Web Services) infrastructure using docker images (https://www.docker.com).

#### Data Processing & Integration

The publicly deployed Ordino instance contains the following data sets (**Supplementary Table S1**):

- The Cancer Genome Atlas (TCGA) https://cancergenome.nih.gov:
  Gene expression, mutation, and copy number data
- Cancer Cell Line Encyclopedia (CCLE) https://portals.broadinstitute.org/ccle [3]:
  Gene expression, mutation, and copy number data
- Project DRIVE [4]: RNAi depletion-screen data (RSA and ATARiS)
- Avana CERES [5]: CRISPR-Cas9 depletion-screen data

##### *TCGA Sample Selection*

All TCGA samples from cBioPortal stage files (downloaded on Sept 29th, 2016) and all samples from the Firehose release 2016-01-28 were included, except for the sample cohorts COADREAD, FPPP, GBMLGG, KIPAN, STES, since these are combined or experimental cohorts.

##### *TCGA Metadata*

The R package TCGAbiolinks (Version 2.5.9) [6] was used to extract sample and patient information for TCGA samples by using a custom-made R script.

##### *TCGA Gene Expression Data*

TCGA data was downloaded from Firehose (https://gdac.broadinstitute.org) using the version `2016_01_28 stddata` between March 11th and March 16th, 2016. Expression data computed by RSEM and represented in tau values were extracted from the archives with the base name `*CANCER.Merge*__illumina*_rnaseqv2__*RSEM_genes__data.Level_3*tar.gz`, where CANCER resolves to all cancer types. Gene expression values were converted to TPM values with the equation TPM = tau * 1e6. Gene names were split into gene identifier and gene symbols. All gene information was mapped to Ensembl 70 gene identifiers using the following procedure: (i) assign all genes for which there is a unique 1:1 correspondence between gene id and gene ids reported by Ensembl, (ii) assign remaining genes if their gene ids and gene symbols map 1:1, and (iii) map remaining genes if their gene ids and symbols map 1:1 after excluding the following gene types: pseudogene, lincRNA, sense. The mapping rate was 19,959/20,531 (97.2%) with respect to mapping genes from the original source. With respect to the samples, only samples of types 1 (Primary solid Tumor), 3 (Primary Blood Derived Cancer - Peripheral Blood), or 11 (Solid Tissue Normal) were kept in the final output. For about 20 samples, more than one aliquot or sequencing result (Illumina HiSeq or GA) was available. In these cases, either the HiSeq sequencing result was taken or the first occurrence in the file in case of a tie situation.

##### *TCGA Mutation Data*

cBioPortal stage files were downloaded on Sept 29th, 2016, and subsequently parsed using a custom-made R script. Additional parsing was necessary, as the transcript for which TCGA reports mutations (column `Transcript_ID`) did not necessarily match the canonical Ensembl transcript. Column "all_effects" of the stage files contains the Variant Effect Predictor (VEP) annotation for additional Ensembl transcripts, which we split on the default delimiter character (";") and inserted into the database. This additional step allowed us to align the mapping between gene symbol and canonical Ensembl transcript ID between CCLE and TCGA.

##### *TCGA Copy Number Data*

TCGA SNP6 copy number segmentation data was downloaded from Firehose (https://gdac.broadinstitute.org) using version `2016_01_28 stddata` between March 11th and March 16th 2016. The segmentation information was obtained from the files `*snp*seg.txt` stored in the archives `*.Merge_snp_*cnv_hg19*.Level_3*gz`.

Gene-wise copy numbers were determined by overlapping the segmentation information with Ensembl 70 gene annotation. If a gene was covered by a single segment, the copy number of the segment was assigned to the gene. If a gene was covered by multiple segments, a weighted average copy number was computed based on the size of the overlap between the gene and each segment.

Relative copy numbers <= 1.0 were considered as "deep deletion", and relative copy numbers >= 3.5 were considered as "amplification".

##### *CCLE Metadata*

Cell line names and descriptions (organ of origin, metastatic site, histology type, morphology, growth type, gender, and age at surgery) were taken from the provider's cell-line data sheet. If a cell line was available from various vendors, the cell-line name was taken from the top rank in a hierarchy of vendors in the following order: atcc, dsmz, ecacc, jcrb, iclc, riken, kclb. A controlled vocabulary of 30 tumor types was derived from the cell-line annotation and assigned to the cell lines with the help of a pathologist.

##### *CCLE Gene Expression Data*

CCLE BAM files were downloaded from https://cghub.ucsc.edu/ using gtdownload in March 2014. BAM files were converted to FASTQ files using SAMtools 0.1.19, bedtools 2.20.1, and the FASTX-Toolkit 0.0.14 as follows: secondary and vendor failed alignments were excluded from the input BAM files, and alignments were shuffled. BAM files were converted to paired-end FASTQ files length-trimmed to 100 bp per read.

All 935 pairs of FASTQ files were then individually aligned to the human genome (hs37d5) using GSNAP version 2012-12-20 together with splice-site information as well as SNP data from the 1000 Genomes Project Phase I (parameters: `gsnap $readgroup -A sam --ordered -n 50 -N 1 -t $threads --gunzip -D $gsnapDB -d $gsnapName -s $spliceiit -V $snpDB -v $snpName --show-refdiff --sam-use-0M $r1 $r2`). Gene and transcript quantification was performed using Cufflinks version 2.0.2 (parameters: `cufflinks -u -p $threads -o $fpkmout --max-bundle-frags 999999999 --no-effective-length-correction --compatible-hits-norm --max-frag-multihits 1 -G $gtf $output.bam`) and the human gene annotation from Ensembl 70. The resulting BAM file with mapped reads was sorted by read names using the SAMtools version 1.0 and used as input to HTSeq version 0.5.3p9 to summarize read counts over gene models (parameters: `samtools view -h -f 0x2 $output.namesorted.bam | htseq-count --quiet --idattr gene_id --type exon -a 20 --stranded no --mode intersection-strict - $GTF > $output.counts`). Various quality control measurements were performed using the following tools: Picard tools version 1.64, RSeQC version 2.3.5, R version 3.0, and FastQC version 0.10.0.

##### *CCLE Mutation Data*

Variant calling of cell lines followed community best practices. The reads were aligned using BWA version 0.5.8 (parameters: `-q 10`) against the reference genome hg19 including decoy sequences (hs37d5), followed by base recalibration and indel realignment and subsequent variant calling using GATK version 1.6. and filtering for artefacts such as polymerase slippage in homopolymer regions, strandness of detected variants, and overall quality. Variants were annotated based on Ensembl 70, and putative germline variation was flagged using external data sets such as dbSNP v135 and data from the 1000 genomes consortium. Putative alignment artefacts were filtered out using a mutation blacklist derived from the Sanger COSMIC Cell line Project VCF files (v70), for which putative artefacts/germline variation is flagged in the VCF files. We computed coverage statistics for each gene in each sample: In the absence of a mutation, we called a gene wild-type if and only if at least 80% of bases of the gene body (excluding the first exon) were sufficiently covered, and NA otherwise.

##### *CCLE Copy Number Data*

SNP6 CEL files were downloaded from https://cghub.ucsc.edu/ in October 2012. Relative copy number segments were computed using the R package `aroma.affymetrix` version 2.13.0 [7-9]: the SNP6 data was processed with the method CRMA v2 followed by CBS segmentation. Afterwards, the copy number segments were overlapped with Ensembl 70 gene annotation analogously to the TCGA processing in order to obtain gene-wise relative copy number values. “Amplification” and “deep deletion” status were also assigned as in the TCGA processing.
Absolute copy number segments were computed using PICNIC version c_release 2010-10-29 [10] with reference files adapted for reference genome hg19 and default parameters. The resulting segments were overlapped with Ensembl 70 gene annotation as in the TCGA processing in order to obtain gene-wise absolute copy number values.

##### *DRIVE Data*

DRIVE (deep RNAi interrogation of viability effects in cancer) is a large shRNA screen of ~8000 genes and ~400 cancer cell lines [4]. Raw data and processed RSA and ATARIS scores were transferred via email by the authors. siRNAs targeting multiple genes were discarded. Gene symbols were translated into Ensembl stable identifiers for genes by using the official gene symbol provided by the Ensembl database Version 70. Cell-line names are identical to CCLE cell-line names and were translated to the Boehringer Ingelheim cell-line nomenclature.

##### *Avana Data*

The Avana single guide RNA (sgRNA) library was used in a large CRISPR/Cas9 loss-of-function screen [5] of ~340 cell lines and ~17,500 genes. Processed CERES scores (representing the estimated gene knockout effects) were taken from the supplement. Gene symbols were translated into Ensembl gene identifiers using the official gene symbol provided by the Ensembl database Version 70. As for the DRIVE data set, CCLE cell-line names were used to translate cell-line identifiers into Boehringer Ingelheim's cell-line names.

#### Case Study 1: Assessment and Selection of Breast Cancer Cell Lines

This case study summarizes an analysis session carried out by a scientist working in a drug discovery team at a pharmaceutical company. In order to identify potential drug targets in a set of tumor types, the analyst performs experiments with cancer cell lines—cultured cells that are derived from tumors and that can proliferate indefinitely in the laboratory. These cell lines are characterized by various properties, such as tumor type (lung cancer, prostate cancer, etc.) and the set of genes that are mutated, deleted, or amplified.

In this case study, the analyst first wants to identify the most relevant amplified genes in the breast cancer cell line HCC1954. Based on these results, the analyst then wants to study a larger set of breast cancer cell lines.

The scientist starts by loading the list of all protein coding genes and adding a column with the relative copy number information for the cell line HCC1954. After sorting by this column, the analyst observes that about 15 genes on chromosome 17 are affected by a large genomic amplification. In order to identify the most relevant gene of these, the analyst adds a column with the gene expression (a measure of activity) in HCC1954 as well as a gene sensitivity score (a measure of importance for cell survival) for HCC1954 (RSA scores obtained from DRIVE data set [4]). The assumption is that amplified cancer genes are highly expressed and that cell lines are sensitive to their removal. Of the highly amplified genes, *ERBB2* (also known as *HER2*) has the highest expression and the most significant sensitivity score, which becomes even more obvious when combining the two columns as stacked bars. It is therefore probably the most relevant gene within this amplified genomic region.

This finding leads the scientist to the question whether *ERBB2* is also highly expressed and frequently amplified in other breast cancer cell lines. To investigate this, the analyst adds a column with the average gene expression, a column with the gene copy number distribution, and a column with the gene amplification frequency across all breast cancer cell lines. He observes that *ERBB2* is amplified in almost 25% of all assessed breast cancer cell lines and that it is often highly expressed.

In order to obtain additional information about this gene, the analyst selects it and opens a series of detail views. Based on the the Expression vs. Copy Number detail view, the analyst notices that there is a clear correlation between copy number and expression of *ERBB2* (the higher the copy number, the higher the expression). The Open Targets detail view provides further information about the gene, for instance, the corresponding protein structure and a list of drugs that target *ERBB2*.

Next, the scientist is interested in obtaining a list of breast cancer cell lines that have both an *ERBB2* amplification and either a *BRCA1* or *BRCA2* mutation (*BRCA1* and *BRCA2* are highly relevant genes in the context of breast cancer). To this end, the analyst opens the Copy Number detail view, filters the cell lines for the tumor type “breast cancer” and sorts the remaining cell lines by their *ERBB2* copy number. After adding columns for *BRCA1* and *BRCA2* mutations, the scientist observes that HCC1954 has the highest *ERBB2* amplification among *BRCA1* mutated cell lines and that HCC1569 has the highest ERBB2 amplification among *BRCA2* mutated cell lines. Finally, in order to obtain further information about these two cell lines, the scientist selects them, opens the COSMIC detail view, and browses the available data.

##### *Analysis Steps and Observations*

*Question: What are the most amplified genes in breast cancer cell line HCC1954?*

- Open list of all protein-coding genes
- Add single cell-line score
  - Cell line: HCC1954
  - Data type: Relative Copy Number
- Sort by copy number column
- Observe: Highest amplification on chromosome 17, affecting about 15 genes

*Question: Which of these genes is the most relevant?*

- Add single cell-line score
  - Cell line: HCC1954
  - Data type: Expression (TPM)
- Add single depletion-screen score
  - Cell line: HCC1954
  - Data type: DRIVE RSA (NB: the lower this value, the more sensitive a cell line is to the depletion of a specific gene)
  - Note that his data is only available for a subset of genes.
- Invert depletion screen score (large bars represent small (very negative) values)
- Observe: Of the highly amplified genes, *ERBB2* (*HER2*) has the highest expression and the lowest sensitivity score. Therefore, it is probably the most relevant gene of this amplicon.
- Combine both score columns to obtain stacked bars
- Observe: Combining the columns highlights the importance of *ERBB2*

*Question: Is ERBB2 also highly expressed and frequently amplified in other breast cancer cell lines?*

- Add aggregated cell-line score
  - Tumor type: breast carcinoma
  - Data type: Expression (TPM)
  - Aggregation: Average
- Add aggregated cell-line score
  - Tumor type: breast carcinoma
  - Data type: Relative Copy Number
  - Aggregation: Boxplot
- Add aggregated cell-line score
  - Tumor type: breast carcinoma
  - Data type: Relative Copy Number
  - Aggregation: Frequency (> 4)

- Observe: *ERBB2* is amplified in almost 25% of all assessed breast cancer cell lines. Further, it is highly expressed.

*Aim: Get more information about ERBB2*

- Select *ERBB2*
- Open Expression vs. Copy Number detail view
- Observe: direct correlation between copy number and expression of *ERBB2*
- Open Open Targets detail view
- Open Pubmed detail view

*Question: In what cell lines is ERBB2 amplified? Select cell lines with ERBB2 amplification that have mutation in BRCA1 or BRCA2.*

- Open Copy Number detail view
- Sort by copy number
- Filter for breast cancer via column menu of column tumor type (also filter out cell lines with unknown tumor type)
- Add single gene score
    - Genes: *BRCA1*, *BRCA2*
    - Data type: AA mutated
- Observe: HCC1954 has the highest *ERBB2* amplification among *BRCA1* mutated cell lines. HCC1569 has the highest *ERBB2* amplification among *BRCA2* mutated cell lines.

*Aim: Show information provided by COSMIC about these two cell lines*

- Select HCC1569 and HCC1954
- Open COSMIC detail view
- Use the drop-down menu to switch between the two cell lines

#### Case Study 2: Prediction of *TP53* Mutation Status

This case study summarizes another analysis session carried out by a research scientist working in a drug discovery team at a pharmaceutical company. In order to identify potential drug targets in a set of tumor types, the analyst performs experiments with cancer cell lines—cultured cells that are derived from tumors and that can proliferate indefinitely in the laboratory. These cell lines are characterized by various properties, such as tumor type (lung cancer, prostate cancer, etc.) and the set of genes that are mutated. One very important gene in the context of cancer is *TP53*. It encodes the p53 protein, whose presence is known to suppress the uncontrolled division of cells. However, when *TP53* is mutated—which is the case in over 50% of cancer patients—it can lose its suppressing function, which results in tumor growth. Due to its important role, scientists want to know whether *TP53* is mutated in a set of cell lines. However, the mutation status of *TP53* is not always known. It has recently been shown that the mean expression level (expression is a measure of the activity of genes) of 13 genes that are biologically related to *TP53* is correlated with its mutation status. The expression level of these genes can therefore be used to predict the mutation status of *TP53* [11]. Furthermore, it has been shown that in many *TP53* non-mutated cell lines, the p53 protein is downregulated through interaction with the protein MDM2. As a consequence, these cell lines are in many cases dependent on the expression of *MDM2*. A downregulation of *MDM2* can result in re-activation of p53 and therefore in the induction of cell death. Hence, cell lines that react sensitively to the removal of *MDM2* are often *TP53* non-mutated, and thus this sensitivity can also be used as a predictor.

In this case study, the analyst first wants to find out how well these two predictors work for the samples contained in the database. Secondly, the analyst wants to predict the *TP53* mutation status for colon carcinoma cell lines for which this information is not available in the database, and seeks further information about the cell lines.

After creating a set containing the 13 *TP53* status prediction genes [11], the analyst loads the list of all TCGA tumor samples, filters for the tumor type colon adenocarcinoma, adds a column showing the *TP53* mutation status, and removes all samples for which the mutation status is unknown. Furthermore, the analyst loads the average expression of the 13 genes. After sorting by the gene expression level, the scientist observes that there is a clear correlation between the gene expression signature and the mutation status of *TP53*: Of the 50 samples with the highest expression only 3 are *TP53* mutated, whereas of the 50 samples with the lowest expression 35 are *TP53* mutated.

The scientist continues the analysis by assessing human cell lines. After loading the list of all available cell lines, the analyst adds columns for the *TP53* mutation status, the average expression of the 13 genes, and the *MDM2* sensitivity data (RSA score from DRIVE data set [4]; a small value indicates high sensitivity). The scientist observes that there is a clear enrichment for *TP53* non-mutated among the cell lines with highest average expression. Furthermore, the *MDM2* RSA values are clearly correlated with the average expression score and the *TP53* mutation status. Based on these observations, the analyst concludes that both predictors are working reasonably well. In order to further improve the *TP53* mutation status prediction, the scientist combines the average expression and the *MDM2* RSA score, observing that the combined score correlates even better with the *TP53* mutation status than the individual scores.

Since the scientist is particularly interested in colon carcinoma, he limits the set of cell lines to this tumor

type and observes that the joint predictor also works well for this subset.
Finally, the analyst wants to predict the *TP53* mutation status for colon carcinoma cell lines for which this information is not available in the database. The scientist includes cell lines for which no *TP53* mutation informations is available. The data set used contains two colon carcinoma cell lines that lack *TP53* mutation status and for which gene expression and DRIVE data is available: MDST8 has a very high combined score and is therefore probably *TP53* non-mutated. NCI-H508 has a very low combined score and is therefore probably *TP53* mutated. In order to assess which genes contribute most to the expression-based *TP53* predictor score, the scientist opens the Expression detail view and observes that the high average expression value in MDST8 can be attributed mainly to the expression of *BAX*, *CDKN1A*, and *RPS27L*. Finally, the scientist opens the COSMIC detail view to obtain further information about the cell lines.

##### *Analysis Steps and Observations*

*Aim: Create gene set for the 13 genes of the expression signature*

- Paste the following list of genes into the gene input field on the welcome page and click “Save”: *AEN*, *BAX*, *CCNG1*, *CDKN1A*, *DDB2*, *FDXR*, *MDM2*, *RPS27L*, *RRM2B*, *SESN1*, *TNFRSF10B*, *XPC*, *ZMAT3*
- Name set *TP53 Predictor*

*Aim: Test applicability of gene signature using TCGA tumor samples*

- Open list of all TCGA tumors
- Filter tumor type *colon adenocarcinoma*
- Add single gene score
    - Gene: *TP53*
    - Data type: AA mutated
- Filter out samples with unknown *TP53* mutation status
- Add aggregated gene score
    - Filter: My Named Sets = *TP53* Predictor
    - Data type: Expression (TPM)
    - Aggregation: Average
    - Compute score only for current sample subset
- Sort by gene expression column
- Observe: There is a clear correlation between gene expression signature and mutation status of *TP53*: Of the 50 samples with the highest expression only 3 are *TP53* mutated, whereas of the 50 samples with the lowest expression 35 are *TP53* mutated.

*Aim: Test applicability of gene signature and MDM2 sensitivity using all cell lines*

- Open list of all cell lines from the start menu
- Add single gene score
    - Gene: *TP53*
    - Data types: AA Mutated and AA Mutation
- Filter out cell lines with unknown *TP53* mutation status

- Add aggregated gene score
  - Filter: My Named Sets = *TP53* Predictor
  - Data type: Expression (TPM)
  - Aggregation: Average
  - Compute scores for all cell lines, not only selected subset (i.e., uncheck option)
- Rename “avg TPM” to “*TP53* Predictor Score”
- Filter out missing values
- Sort by TP53 predictor score
- Observe: There is a clear enrichment of *TP53* non-mutated among the cell lines with high score.
- Add single depletion-screen score
  - Gene: *MDM2*
  - Data type: DRIVE RSA (NB: the lower this value, the more sensitive a cell line is to the depletion of the gene of interest)
- Invert score so that large bars represent small (very negative) values
- Filter out missing values
- Sort by *MDM2* RSA score
- Observe: Small *MDM2* RSA values (large bars) are correlated to the expression score (*TP53* predictor score) and the *TP53* mutation status
- Combine the two score columns by dragging one column onto the other, which results in a weighted sum column
- Sort by the combined column
- Observe: Combined score correlates even better with *TP53* mutation status than individual scores.

*Aim: Assess predictors in colon carcinoma cell lines*

- Filter for tumor type *colon carcinoma*
- Observe: Combined score correlates well with *TP53* mutation status for colon carcinoma cell lines.
- Conclusion: The TP53 target score as well as the *MDM2* RSA score can be used to predict a *TP53* mutation status of cell lines for which it is not available.

*Aim: Predict TP53 mutation status for colon carcinoma cell lines for which this information is not available in the database and assess which genes contribute most to the TP53 predictor score. Furthermore, obtain additional information from COSMIC about these cell lines.*

- Include cell lines with unknown *TP53* mutation status.
- Observe: The data set used contains two colon carcinoma cell lines that lack *TP53* mutation status and for which gene expression and DRIVE data is available: MDST8 has a very high combined score and is therefore probably *TP53* non-mutated. NCI-H508, however, has a very low combined score and is therefore probably *TP53* mutated.
- Select the two cell lines and open Expression detail view
- Observe: The high average expression value in MDST8 is caused mainly by the expression of *BAX*, *CDKN1A*, and *RPS27L*
- Open COSMIC detail view and browse available information
